# A flexible fluid delivery system for rodent behavior experiments

**DOI:** 10.1101/2024.04.27.590287

**Authors:** Bruno F. Cruz, Paulo Carriço, Luís Teixeira, Sofia Freitas, Filipe Mendes, Dario Bento, Artur Silva

## Abstract

Experimental behavioral neuroscience relies on the ability to deliver precise amounts of liquid volumes to animal subjects. Among others, it allows the progressive shaping of behavior through successive, automated, reinforcement, thus allowing training in more demanding behavioral tasks and the manipulation of variables that underlie the decision making process (*e*.*g*.: reward magnitude). Here we introduce a stepper-motor-based, fully integrated, open-source solution, that allows the reproducible delivery of small (<1 *µL*) liquid volumes. The system can be controlled via software using the Harp protocol (*e*.*g*.: from Bonsai or Python interfaces), or directly through a low-level I/O interface. Both the control software and electronics are compatible with a wide variety of motor models and mechanical designs. However, we also provide schematics, and step-by-step assembly instructions, for the mechanical design used and characterized in this manuscript. We provide benchmarks of the full integrated system using a computer-vision method capable of measuring across-trial delivery of small volumes, an important metric when having behavior experiments in mind. Finally, we provide experimental validation of our system by employing it in a psychophysics rodent task, and during electrophysiological recordings.

## Introduction

Modern experimental neuroscience research relies on training animals in tasks that isolate or exacerbate demands for specific cognitive variables of interest [Gomez-Marin et al., 2014]. Given the non-verbal nature of most subjects, stable performance in such paradigms is often achieved through the successive reinforcement of simpler behaviors, a process known as “shaping” [Jones and Skinner, 1939]. These reinforcers may take several forms. Nevertheless, for both historical and practical reasons, liquid rewards have become a staple in most widely studied animal models [Guo et al., 2014].

Due to their simplicity and compactness, gravity-based passive systems, most commonly implemented using valves, are widely adopted by the neuroscience community. With this approach, the volume of the delivered fluid is determined by the duration a valve remains open. Despite their convenience, gravity-based systems routinely face operational issues. First, due to changes in the fluid resistance (e.g.: biofilm growth in tubing), calibration values tend to drift across days, requiring frequent maintenance and re-calibration. Second, the relationship between reward amount and valve opening time is often non-linear, especially for small volumes, requiring calibration over several reward sizes. Finally, since the fluid flow rate is constant under a given value of hydraulic pressure, it is challenging to decouple delivered liquid volume from the total delivery time.

Alternative active systems, such as syringe pumps, have the potential to solve all the aforementioned problems. Unfortunately, currently available commercial systems are prohibitively expensive, often lack a flexible control system to fully satisfy the experimental needs of the users, and are difficult to implement at scale.

Here, we present and characterize an open-source syringe pump system (Figure 1) with scalable neuroscience experiments in mind. We provide detailed instructions and a parts-list that allow the system to be fully assembled using widely available off-the-shelf and custom 3D printed parts. In addition to mechanical designs, we also developed a flexible control system that affords a large range of customizability over the system’s function. This control is implemented in a custom-designed printed circuit board (PCB) that implements the Harp protocol.

**Figure 1.**
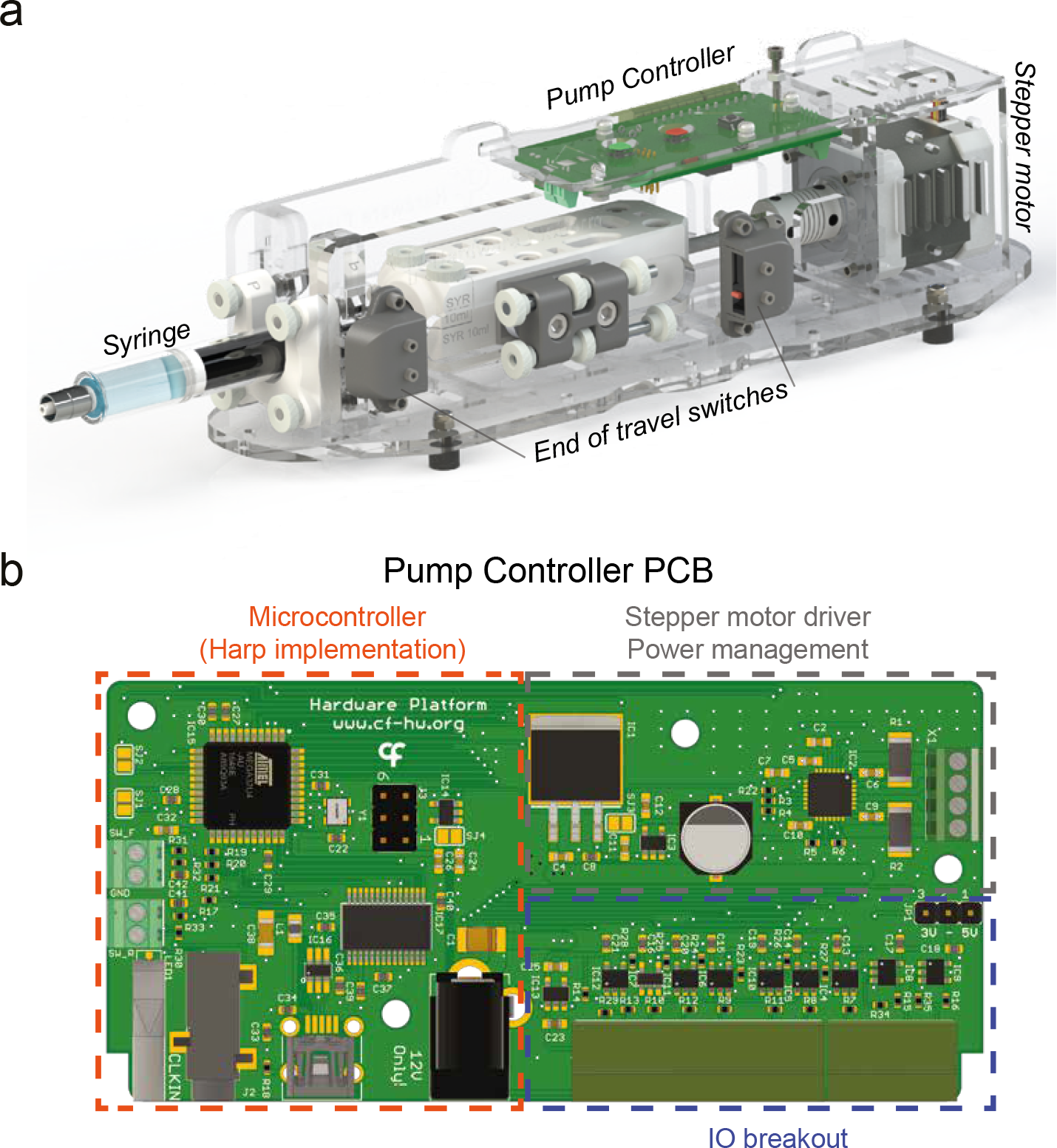
Syringe pump system. (**a**) 3D model of the fully assembled syringe pump system. Controller PCB, syringe, switches, and stepper motor are highlighted. (**b**) Diagram of Controller PCB. The three main sections of the board are highlighted: Microcontroller, which implements the Harp protocol. Motor driver and power, which provide the low-level logic to drive the stepper motor, and the I/O breakout, that affords users with input and output lines which can be used to control and monitor the function of the system, respectively. See Methods for further details

Additionally, the provided designs allow users to control the pump in a variety of ways. From triggering pre-defined protocols with a single square pulse to fully specify the behavior of the pump using the Harp Protocol. The latter interface, affords communication with Bonsai [Lopes et al., 2015], which in turn integrates the device in an ever increasing ecosystem of software for experimental behavior control and acquisition.

Similarly to other open-source systems [Wijnen et al., 2014, Amarante et al., 2019], we characterized the error associated with the delivery of large volumes. Additionally, since rodent experiments often rely on the delivery of microlitre range rewards we also designed a simple assay, leveraging computer vision, to characterize the performance of the pump in single-bolus events.

To validate the usefulness of the system, we varied the amount of reward delivered and show that this manipulation can quickly and reversibly alter rat’s choice behavior. Finally, we highlight the high compatibility of the described syringe pump with electrophysiology recordings, and demonstrate no detectable electric artifact was observed.

## Results

### Characterization of sub-microliter volume delivery

Given the common reliance of experimental neuroscience on the delivery of small liquid volumes, we sought to first benchmark how our system performs in this regime. Single small-volume events are experimentally challenging to accurately measure. Other systems have relied on inferring the average predicted volume of a single bolus, from repeated measures. Despite straightforward, such method cannot resolve trial-to-trial variability. With this in mind, we developed a quantitative computer-vision-based assay, that can provide a metric of such variability (Figure 2). Our methods relies on the videography of a small-diameter glass capillary with well-characterized and reproducible dimensions. By adding small amounts of food colouring to the fluid, we can reliably segment the area of the glass filled with liquid. Assuming *Area*_*pixels*_ ∝ *V olume*_*mL*_, we can use it as a linear proxy for delivered volume.

**Figure 2.**
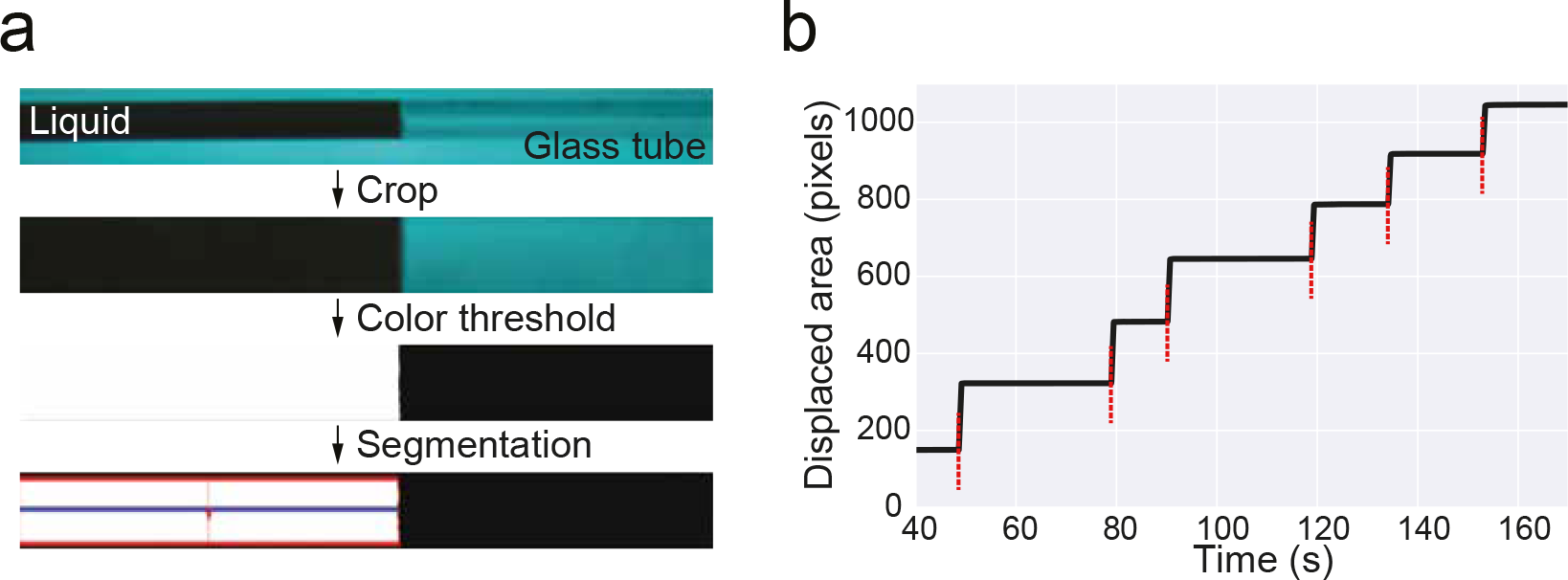
Tracking small single-bolus events. (**a**) Schematic of the computer vision algorithm for measuring small microliter-range volumes. From top to bottom: cropped, thresholded, and segmented the area of the capillary filled with liquid. Area was taken as a proxy of delivered volume. (See Methods for further details) (**b**) Example trace of the measured area in A), as a function of time during one of the experiments. Red vertical dashed lines represent liquid delivery events.

Using this approach, we picked four distinct liquid volumes, in a range relevant to rodent behavior experiments (0.33 *µL* - 10 *µL*). Based on the mechanical specifications of our system and mounted syringe models, we calculated the number of steps expected to achieve such volumes. Next, using the videography data, we constructed a “protocol-onset triggered” trace that can be used to quantify single-trial total delivered volume (during the stable phase of the trace), as well as its fluid kinetics (Figure 3-a). This procedure provides an estimate of variability across single delivery protocols, on top of revealing a close linear relationship between theoretically predicted and delivered volume. This result held true not only across different setups (*Pump A vs. Pump B*), but also across mounted syringes with different dimensions (10 *mL vs*. 5 *mL*) (Figure 3-b,c). Finally, while the spread of single trial delivered volumes grew with theoretically calculated values, its coefficient of variation remained relatively stable in all conditions (Figure 3-d,e).

**Figure 3.**
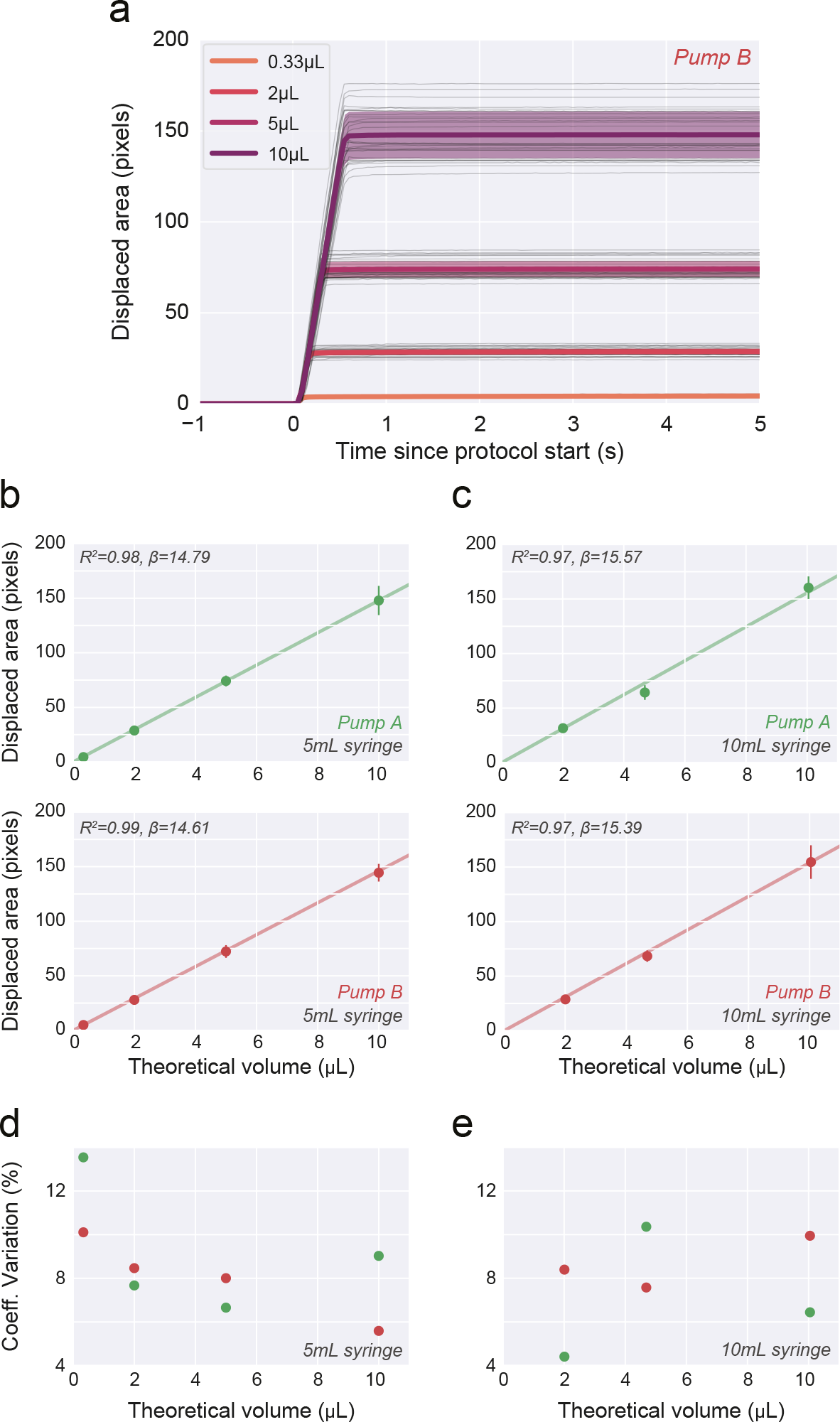
Single-bolus protocol calibration. (**a**) Time course of displaced area aligned on protocol onset (*t* = 0) for four distinct theoretically expected delivered volumes. Thin and thick lines correspond to single trials, and averages for a given expected volume, respectively. Shaded area depicts s.t.d. (**b, c**) Total displaced area for the four protocols. Each point shows mean*±*s.t.d.. across 30 replicates, per volume, in two different pumps (*Pump A* and *Pump B*, green and red, respectively) and using two different glass syringe sizes (5 mL and 10 mL, (**b**) and (**c**), respectively). (**d, e**) Coefficient of variation (s.t.d./mean) calculated from the data shown in (**b**) and (**c**), respectively.

### Flow-rate modulation

Our characterization method also affords the chance of measuring its single-trial time-course dynamics (*i*.*e*. flow-rate, pixels *s*^−1^). Theoretically, flow rate should be a function of the number of steps the driver takes per unit time. As a proof-of-concept, we performed an experiment where we varied the time between each train of pulses (protocol) and characterized how these changes affected the small amounts of liquid delivered per unit time (Figure 4). Our results showed that, as predicted, the displaced area in the capillary per unit time monotonically decreases with larger intervals between protocols. While this relationship is clearly not linear, the reproducibility of such dynamics across trials affords the chance of precisely calibrating the flow-rate for a specific system.

**Figure 4.**
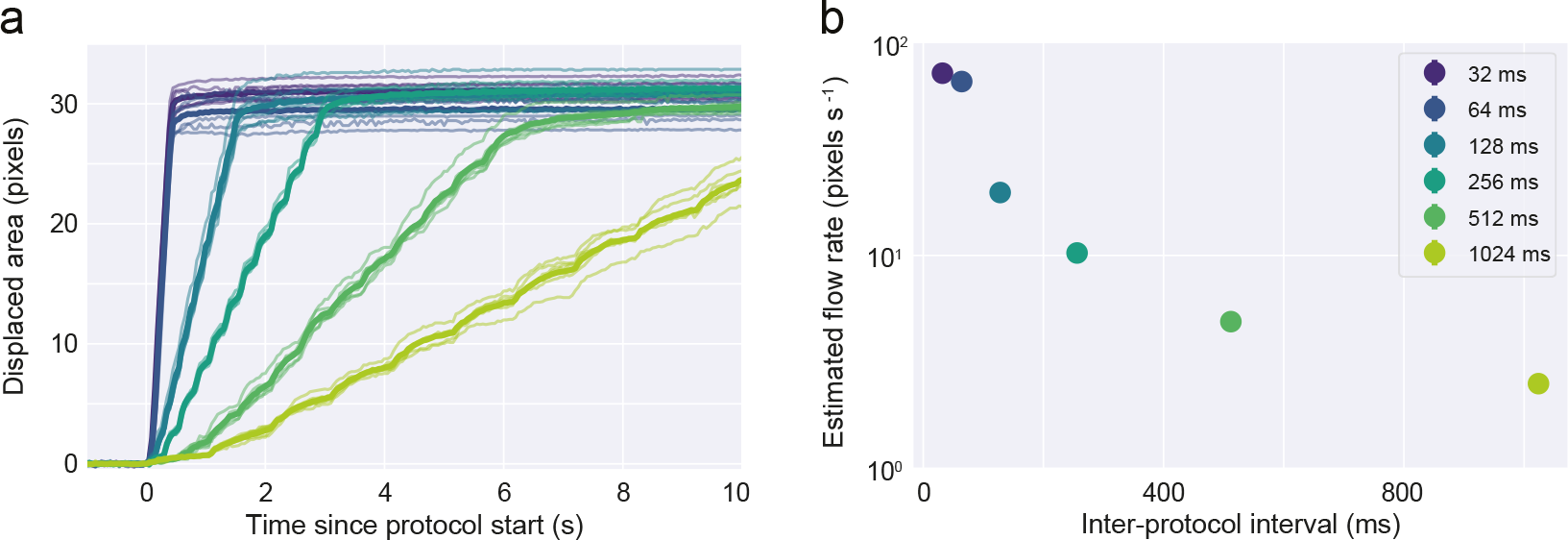
Syringe pump affords dynamic control over the flow rate. (**a**) Time course of delivered volume (displaced pixel area) aligned on protocol onset (*t* = 0) for different inter-protocol-interval values (IPI). Thin and thick lines represent single trials, and averages, for a given IPI, respectively. (**b**). Estimated flow rate (pixels s^−1^) for all tested IPI (mean*±*s.t.d.., n = 6 trials for each IPI).

### Calibration of large volumes

After characterizing the small-bolus delivery capabilities of the system, we decided to follow a typical, and more experimentally amenable, calibration protocol. Measuring single small-volume delivery events presents a technical challenge in that equipment capable of reliably measuring such small amounts is rarely available. As a result, experimenters rely on measuring the outcome of several hundred repeats of the same protocol. While this strategy effectively blinds the researcher to inter-protocol variability, it allows the measurement of larger volumes, with smaller systematic errors, and thus the estimation of an average single-protocol delivery volume. Following such a protocol (see Methods), we demonstrate how calibration curves can be obtained from the setup, as well as validate the theoretically predicted amount of delivered volume calculated from the physical specifications of the system (Figure 5). As predicted, both the quality (*R*^2^), and the slope (*β*), of the linear fit highlight the high-quality match between expected and measured volumes (Figure 5a). Despite not being stable throughout the full range of probed delivered volumes, the coefficient of variation that results from measurements of this method is strikingly low (Figure 5b).

**Figure 5.**
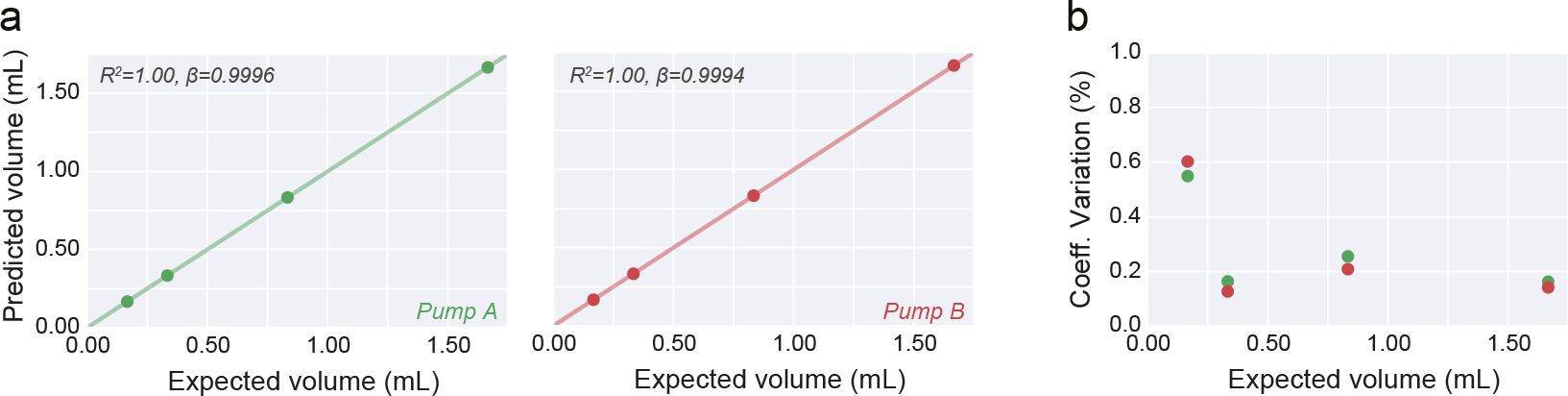
Large volume protocol calibration. (**a**) Delivered volume (mean*±*s.t.d.) across 20 performed delivery protocols for 4 distinct large volume amounts, in two independent pump systems (*Pump A* and *Pump B*, green and red, respectively). “Predicted volume” was calculated by weighting the delivered liquid and assuming a water density of 1 g/mL. “Expected Volume” was calculated using the theoretically displaced syringe volume. *R*^2^ and slope (*β*) resulting from the linear fit to the data are shown in the top-left corner.(**b**) Coefficient of variation (s.t.d./mean) calculated from the data shown in (**a**.

### Calibration stability and comparison against gravity-based systems

In the previous section, we show how calibration of the system can be achieved in an experimentally amenable fashion. Whilst such a procedure should take no more than a few minutes, it can quickly become unfeasible if several water delivery systems are routinely used simultaneously. As a result, we next benchmarked how stable the calibration is by performing longitudinal, weekly, calibration procedures. Our data shows that, at least, up to one month after the initial procedure (*Day 0*), calibration curves are identical (Figure 6a, left).

**Figure 6.**
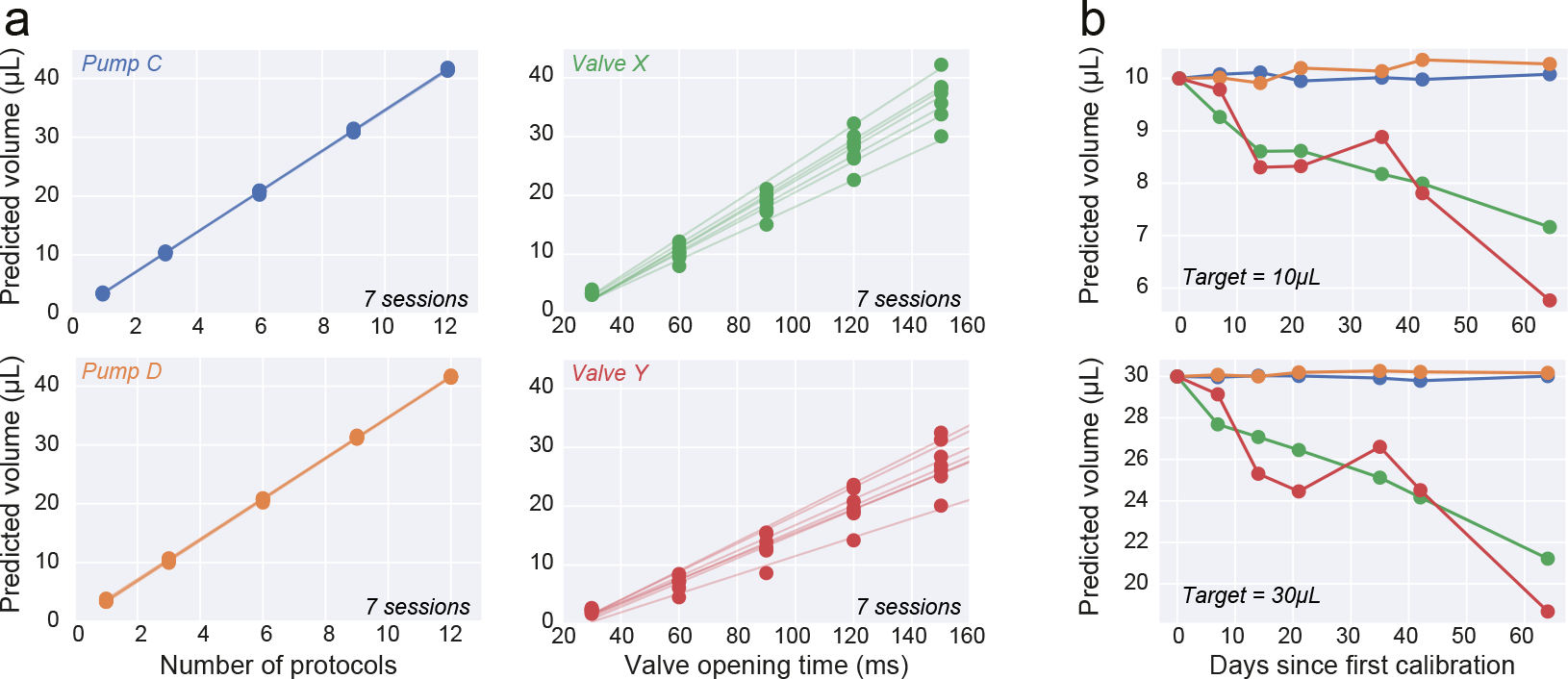
Comparison of calibration stability between the gravity-based solution and the presented pump system. (**a**) For each system, the calibration (See Methods) consisted of measurements of delivered liquid as a function of the number of pre-programmed delivery protocols, or valve opening time, for the *Pump* (left) and *Valve* (right) systems, respectively. Individual fits correspond to a single-day calibration protocol. For each system, we tested two devices independently (distinct colours). (**b**) Predicted delivery volume for two arbitrary volumes (10 *µ*L and 30 *µ*L, top and bottom, respectively) based on the calibration linear fits derived from A). For each day, the fit calculated from day 0 was used as to infer that value would have been delivered had the calibration curve remained the same.

Next, we considered the most common alternative to a syringe-pump system, in experimental behavior neuroscience: gravity-based systems. These systems rely on a liquid reservoir at a higher potential than the system’s outlet. While simpler and generally more affordable, these systems are usually subject to changes in the fluid path resistance that lead to systematic differences in delivered volume over time. This problem can be partially circumvented by investing time on regular setup calibration and maintenance, which tends to hinder the scalability of this solution.

To compare the stability of the presented syringe pump system versus a common gravity-based system, we decided to perform and track regular calibration results of both the systems Figure 6.

A calibration procedure identical to the one described in the last section was followed. As expected, both systems show daily calibration curves consistent with a linear behavior (Figure 6a). However a clear change in slope over days can be observed in both valves (Figure 6a). Using the daily calibration values, one can ask: “how much liquid volume would be delivered, had the experimenter not performed the calibration procedure, and kept using the values from Day 0” (Figure 6b). This analysis revealed that gravity-based valve systems steadily drift over days, whereas syringe pumps remained stable. The general decrease in delivered volume is likely due to a drop in flow rate due to the accumulation of biofilm in the tube and valves.

### Experimental use cases

We next show two examples of the presented syringe pump system use in a neuroscience laboratory setting. Example experiments highlight both the day-to-day easy of operation and also the compatibility of the system with other methods.

In a first example, we tested rats in a two-arm bandit task, where delivered reward amount was parametrically varied using our system. On each block (see Methods), the amount of reward at each arm was randomly drawn from a set of five possible values. Both tested animals (N=2) appeared to be sensitive to the relative amount of reward delivered and showed a systematic preference towards the highest-reward amount option, in a manner consistent with its relative difference in magnitude (Figure 7). This preference quickly reversed at block transition, modulated suggesting that animals did not need to rely on several trials to determine the highest-value arm.

**Figure 7.**
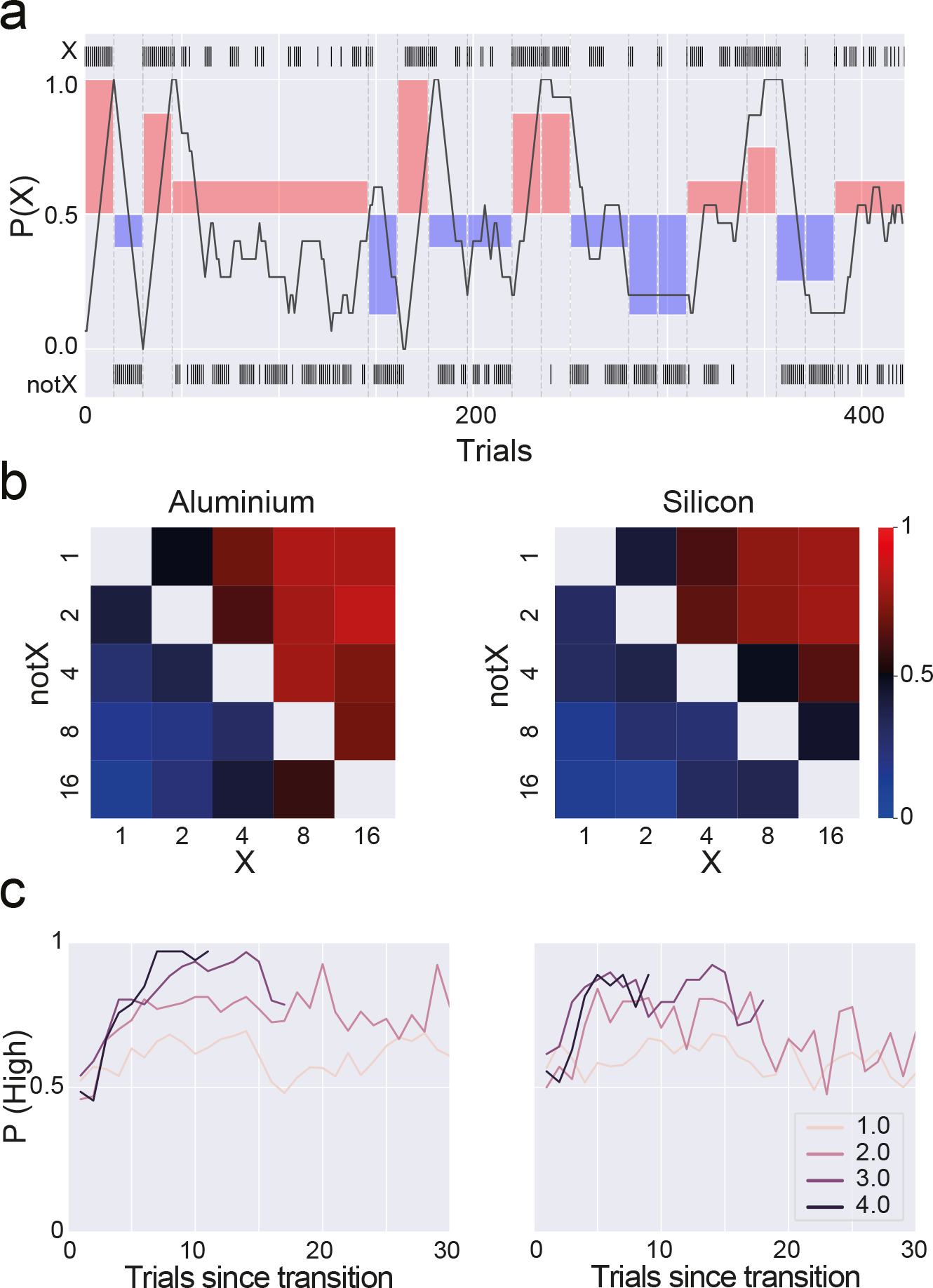
Rats quickly reverse choice preference on delivered liquid reward amount. (**a**) Example session. Animals alternate between the two available choices (“X” and “notX”, top and bottom raster plots, respectively) in a way that appears consistent with the highest-value choice (shaded area). Thick dark line depicts the probability of choosing “X” in a rolling window of 15 trials. (**b**) Probability of choosing reward giving nose-port X as a function of the possible reward amount combinations given by each nose-port. Values in the heatmap axes are the number of protocols given on each reward delivery, by each reward nose-port (X and notX). (**c**) Probability of choosing the highest rewarded side over the trials following block transition. Lines correspond to the absolute value of the difference between rewards at each nose-port in logarithmic units (base 2). The highest the difference between the two rewards available on a particular block, the sooner animals converge to the highest rewarded nose-port, suggesting the need for fewer reward samples (trials) as the difference between rewards increases.

Finally, given the current reliance of the Neuroscience field on electrophysiological recordings, it is pertinent to ascertain the compatibility of the described system with such techniques. Previous reports of induced artifacts stemming from stepper-motor operation near these systems [Amarante et al., 2019]. As a result, we performed acute electrophysiological recordings from a mouse’s brain (Methods) while simultaneously operating the syringe pump (Figure 8). We leveraged one of the digital outputs in the board to send a pulse on each “STEP” instruction, thus providing precise sub-millisecond alignment to the activity traces. Our data showed a clear absence of artifacts aligned to, or surrounding, each pulse, while still preserving easily identifiable single-unit spiking activity, thereby validating the use of the presented syringe pump system during electrophysiological recordings.

**Figure 8.**
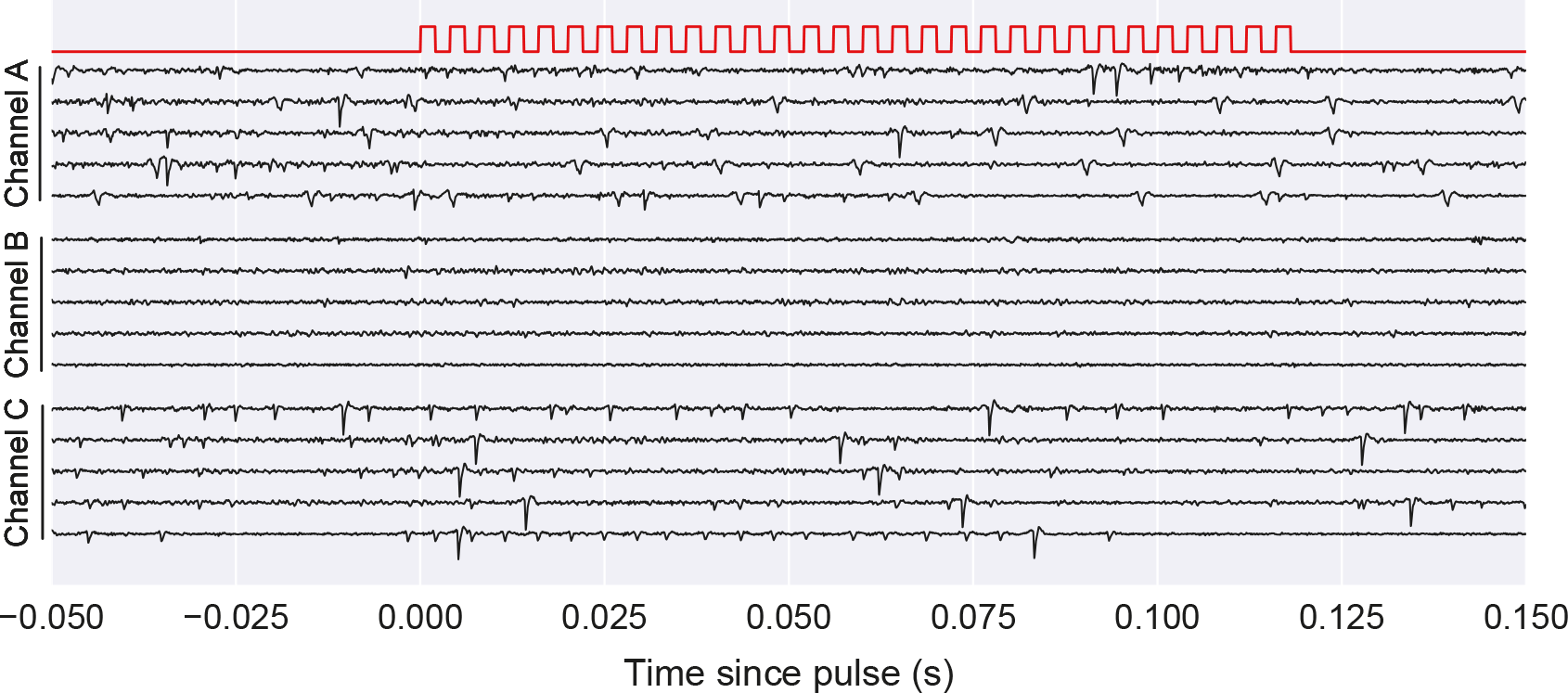
Compatibility with electrophysiological recordings. Example single trial traces (four trials) for three simultaneously recorded channels (A-C) aligned to the start of a protocol of several pulses (red trace). See Methods for further details. Notice the presence of spiking activity in Channels A and B, along with the lack of any clearly identified stepper-motor-induced electrical artifact.

## Discussion

Inspired by previous projects ([Amarante et al., 2019, Wijnen et al., 2014]), we present a fully integrated, and characterized, system for micro-litter range fluid delivery targeted towards neuroscience behavior experiments. We designed a simple protocol which allowed us provide an extensive characterization of trial-to-trial variability in the fluid delivery dynamics. An import stack of tests when considering the intended use of this system for rodent behavior experiments.

We provide an alternative to gravity-based passive systems, by designing a scalable and affordable solution that allows for dynamic control over reward delivery, without suffering from the long-term stability issues often associated with the former systems. Additionally, we also highlight the afforded control gained over flow-rate that such an non-passive system allows. Importantly, we also tested the compatibility of the system with common neuroscience experimental demands by deploying it during rodent behavior and electrophysiological recording experiments. Finally, we provide users with several options to interface with the system, affording flexible configuration and easy integration into already-existing experimental rigs.

## Methods

### Syringe Pump construction

The presented syringe pump system is composed of a mechanical assembly, an electronic controller board, and an accompanying firmware/software stack that affords different levels of control over the system’s behavior. All resources (interface software, firmware, and mechanical designs) are available from https://github.com/harp-tech/device.syringepump.

#### Mechanical construction

The mechanical assembly of the syringe pump was designed to be easily produced and assembled without any particular expertise. All parts necessary for the build are included in the bill of materials. Most parts are readily available off the shelf. Additionally, custom structure parts can be in-house laser-cut and 3D printed, if such resources are available, or sourced from third-party manufacturers, using the provided designs. In addition to the bill of materials, we also documented the assembly process with step-by-step instructions, including photographs, that should be followed to guarantee the best possible assembly of the structure.

In the provided mechanical configuration, we opted for a NEMA 17 Stepper motor (HT17-275 from Applied Motion) due to its high precision and torque rating. Specifically, the combination of motor step resolution (1.8 deg/step) together with a 0.8 mm pitch length driving rod leads to a theoretical linear resolution of 4 *µm*/step, which can be further increased by using step multipliers of up to 1/16. Additionally, to characterize the system, we chose two glass syringes models with total volumes compatible with those routinely used for animal experiments (5 and 10 *mL*, 1000 Series Syringes, Hamilton). Taking their cross-section diameter into consideration, the final single-full-step theoretical resolution will be 0.33 and 0.66 *µL/step* for the 5 and 10 *mL* syringe models, respectively.

It should be noted, however, that the present system will be able to accommodate a large range of other options (be it stepper motors or syringes), with little to no modifications to the suggested design.

### Electronics

The syringe pump is controlled by a custom-made printed circuit board which is comprised of three main blocks: a microprocessor, an interface logic circuitry and, a motor driver (Figure 1).

The microprocessor block consists of an ATxmega microprocessor (ATXMEGA128A4U-AU, Microchip Technology) that implements the Harp protocol https://github.com/harp-tech/protocol, a USB to serial UART interface and a stereo phone jack for temporal synchronization between Harp devices. The Harp protocol allows the syringe pump to be flexibly controlled externally through the USB interface with any host implementing the Harp API (*e*.*g*.: via Bonsai[Lopes et al., 2015] or Python).

The interface circuitry block consists of logic buffers that provide a direct low-level interface with the micro-stepper driver (A4988SETTR-T, Allegro Microsystems) to the user (bypassing the microcontroller block). Moreover, it provides access to the I/O breakout that can be used to trigger protocol delivery with other external, TTL-compatible, devices, supporting input voltages up to 5 V. The voltage of the digital outputs can also be configured in the board through a jumper to select between 3.3 V and 5 V voltage output logic.

The modularity of the board design affords users the option to assemble a simpler version without the microcontroller block, for applications wherein low-level control is sufficient. This significantly cheaper version is assembled using the same PCB schematic, with minimal changes to the board components, and can be found in an alternative bill of materials provided.

Despite having been designed for the herein described configuration, the design of the control system (including the Harp API) is compatible with other types of bipolar stepper motors (assuming an output drive capacity of up to 12 V and ±1.7 A).

### Syringe pump user control

We designed the syringe pump system with three distinct interface levels that afford experimenters control over the pump according to their experimental needs. In addition to (Figure 9), we offer a brief description of the three available options:

**Figure 9.**
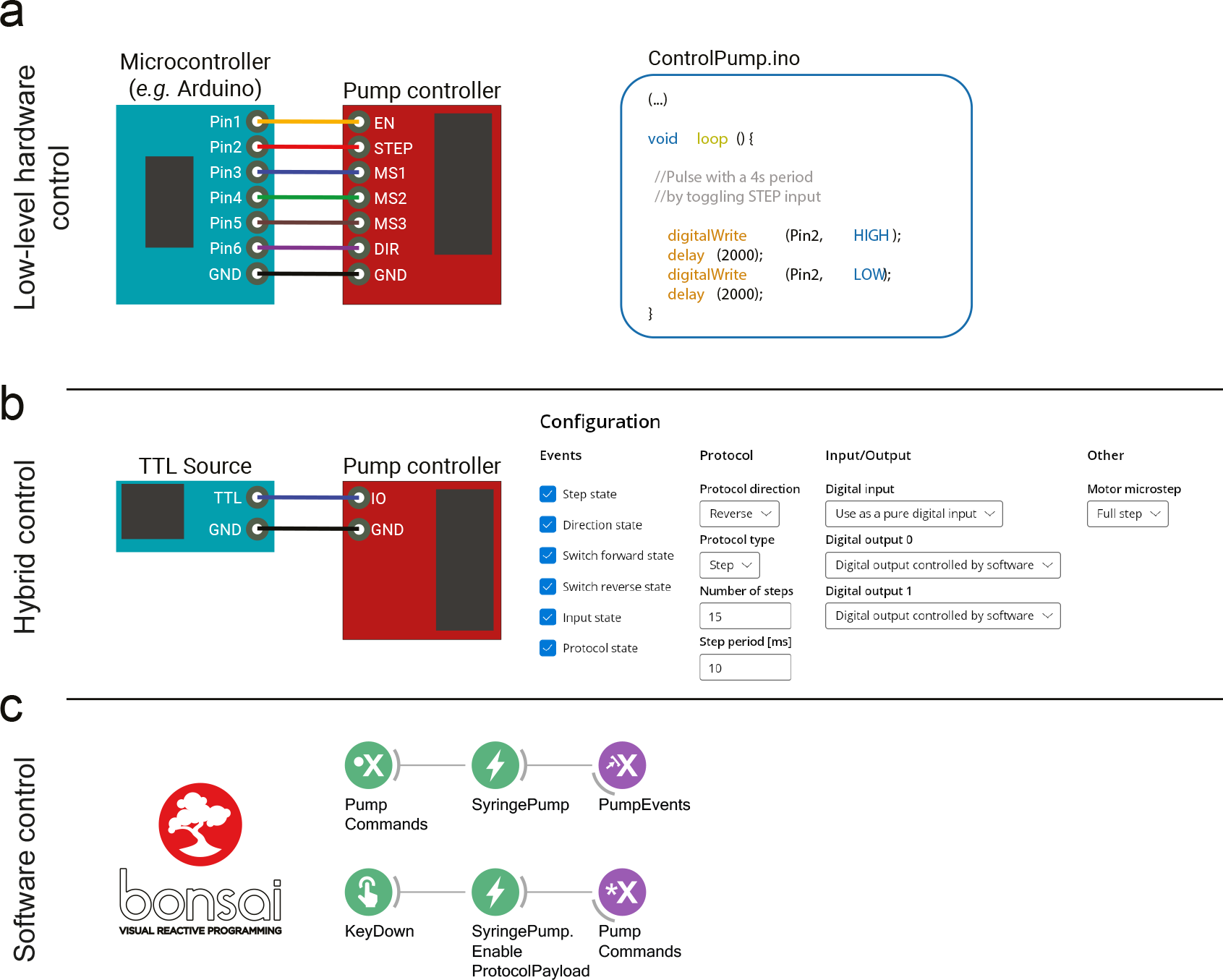
The current system affords three distinct levels of interface. (**a**) Low-level hardware control expects the user to fully define the control logic for the stepper motor driver. This can be achieved by implementing such logic in a microcontroller (*e*.*g*.: Arduino, right textbox) that defines the state of all input control pins to the stepper-motor driver. It should be noted that this mode does not require the full PCB to be populated, since it does not rely on the Harp core protocol implementation. (**b**) Trigger-based control allows the user to use an external trigger event (via transistor-transistor logic, TTL) to playback a pre-defined delivery protocol. The default protocol values (*e*.*g*.: volume and flow-rate) can be modified using a provided graphical-user interface, or Bonsai. (**c**) Software control allows the user to fully parameterize, and trigger, protocols from a computer host running Bonsai without the need for any external hardware triggers. Sub-millisecond synchronization can be achieved via Harp protocol, or by a configurable digital output event.

- Low-level control - This option relies on directly controlling the I/O interface of the micro-stepper driver. This is achieved by using an external source (*e*.*g*.: Arduino microcontroller) to generate the necessary input logic. Eight inputs lines are exposed: GND (ground), EN (enables the driver when low), MS1-3 (determines the step-size of the motor, 1 to 1/16 times a full step), DIR (determines the direction of rotation) and STEP (triggers a micro-step). It should be noted that this control logic is common and documentation is extensively available. Additionally, the PCB exposes two outputs that correspond to “end-of-travel” switches, that can be used to implement operation safety-stops. This control option does not require the full board to be assembled, as it does not rely on the microcontroller block for control and, while it affords the user the most flexibility, it requires an external source of logic control that might be cumbersome to set up and unnecessary for the vast majority of applications. Thus, the next two options abstract some of the low-level control away from the user at the cost of some flexibility.
- Trigger-based Control - This option allows the user to configure a *“protocol”* that is triggered with the detection of a TTL rising edge. This control logic is extensively used across many platforms (*e*.*g*.: Med associates, Open-Ephys, Arduino) making the system easily integrated with existing experimental rigs. The protocol can be configured via a custom-designed graphic user interface (Figure 9) that leverages the Harp protocol to configure the microcontroller. Available settings currently include an option to run in “Step Mode”, wherein the user sets number, duration, and frequency of the steps per protocol). The behavior of the I/O pins can be configured via this GUI to provide users with hardware events that can be used to interface with third-party devices (e.g.: see Figure 8) It should be noted that, contrary to the previous mode, “end-of-travel” events (*i*.*e*., when the limit switches are triggered) are automatically handled by the controller, to prevent accidental mechanical damage to the system.
- Software Control - The third option allows the user to control the syringe pump system through a software API. This option can be seen as an extension of the last, with the protocol being triggered by a software, instead of an hardware event. This option is available to any host running the Harp API, namely Bonsai. Bonsai is a visual programming language that specializes in asynchronous data acquisition and has seen growing adoption from the neuroscience community. Users can change an large number of settings (identical to the “triggered control” version) and trigger a protocol on any Bonsai event. As an example, the user can periodically deliver a constant amount of reward (Figure 9), dynamically change the amount of liquid delivered on every trial, contingent on some behavior/physiological event, or trigger the delivery of water aligned to when the animal enters a given area of an arena. Finally, the Harp protocol allows the user access to hardware timestamped events (*e*.*g*.: protocol onset), allowing for easy event alignment.

### Calibration

#### Large volume calibration

The pump is compatible with a wide variety of syringes. However, it is advisable to calibrate the full system to account for potentially significant differences across parts. Small volumes of delivered liquid are technically challenging to measure, as a result, calibrating liquid delivery systems is usually performed by repeating the same protocol a large number of times. This strategy rests on the assumption that single steps are relatively reproducible and the mean across several dozens to hundreds of repetitions is thus representative. We calibrated the system by varying the number of steps per protocol, which was then repeated 200 times, waiting 250 ms between the end of each protocol and the start of the next. We then weighted the amount of collected liquid to yield the total collected volume (we assumed a water density of 1 g/mL at room temperature). Single protocol volume was thus given by 1:

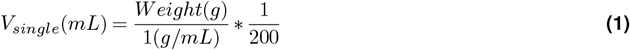

To assess reproducibility across runs, we repeated this protocol twenty times. Depending on the experiment, we also adjusted the range of calibration values. Critically, all tested volumes laid on top of a linear calibration regime, and outcomes were consistent with the predicted theoretical volumes (Eq. S (1)).

We followed protocol identical to the syringe pump system to calibrate the solenoid valves (LHDA1231215H, Lee Co.). Briefly, we calibrated the valves by setting the water reservoir (a 20 mL plastic syringe) at a height identical to what we routinely use in our laboratory for behavioral experiments (∼30 cm). We systematically varied the opening time (between 10 and 150 ms) to yield different delivered volumes. Both the number and time between trials were identical to the ones used in the pump systems. Hardware control logic was implemented in an Arduino Microcontroller (Arduino Mega, Arduino).

### Single bolus calibration

While technically amenable, measuring the outcome of several hundred protocols might obscure variability across single trial events. We thus developed a simple computer-vision-based method to measure small volumes. Briefly, the end of a syringe was fitted with a plastic adapter with a small glass capillary (Model 211713, Vitrex) glued to the end (Figure 1). A computer vision camera (Model FL3-U3-13S2C-CS, FLIR) was used to image the capillary. In order to increase the contrast between liquid and background, we diluted a small amount of red colourant (Erythrosin B, Sigma-Aldrich) that, when combined with LED illumination, resulted in the image shown in (Figure 2).

The video acquisition and processing routine were implemented in Bonsai. To measure the amount of displaced liquid inside the capillary, we defined a region of interest, excluding optical artifacts resulting from glass-induced diffraction, and binarized the image thresholding the pixel values in RGB colour space. We then used this binarized image to segment the area corresponding to the displaced liquid volume. We choose the horizontal axis of the region as our reported metric. Data were acquired at 10 frames-per-second and synchronization was achieved by sending a short TTL pulse to the camera each time a protocol was started. Syringe pump control logic was implemented using the low-level control mode of the system and an Arduino microcontroller.

Unless otherwise stated, the inter-pulse-interval was set to 4 milliseconds. Before each run, up to 5 small pulses were applied to ensure that every protocol started from comparable conditions. These pulses were discarded from all analyses. The interval between each protocol was drawn from a uniform distribution (10 to 20 s). Due to the size of the capillary, some volume values required cleaning the tube between trials. We cleaned it by flushing ethanol and air-drying it.

For the dynamic flow rate experiment shown in Figure 4, the inter-pulse-interval was systematically varied in order to control the output flow rate. We estimate flow rate by linearly fitting, on each trial, the displaced area to time (between protocol onset and the stabilized epoch).

### Experimental validation

#### Animal behavior experiments

Two Long Evans, 5 month-old, male rats were trained on a variant of a Two-armed bandits task [Ito and Doya, 2009] to assess reward preference. Briefly, subjects were placed in an experimental box with access to four nose ports, two for initiation, and two for reward delivery. Reward volumes were drawn from a pre-defined set of calibrated values (3.46, 6.92, 13.84, 27.68, and 55.36 *µ*L) and delivered using the syringe pump system. A light on top of the initiation ports signalled trial availability. After initiating, animals were required to fixate at the centre nose port for a short period of 100 milliseconds, after which they would be allowed to collect a reward at one of the two side-reward ports. Importantly, the amount of water delivered at any port was constant within a given session block, but always different across the two ports. After reaching a preference criteria of >80% towards the highest reward nose-port over the last 15 trials, a block transition would be triggered, the trial initiation switched to the opposite nose port (*e*.*g*.: odd and even numbered blocks would be initiated at the north and south nose ports, respectively). At block transitions, we forced reward amounts on each nose-port to be different, but otherwise allowed for any other combination. In particular, the highest rewarded nose-port need not to change between blocks.

#### Compatibility with electrophysiological recordings

Previous studies report induced electrical artifacts from some syringe pump systems [Amarante et al., 2019]. We thus tested the compatibility of our system with electrophysiological techniques by recording these signals while triggering the micro-stepper driver in close proximity. We followed the protocol for acute recordings previously described in [Cruz et al., 2022]. In short, mice (8-10 weeks old) were anaesthetized with Isoflurane (1-2 % at 0.8 L/min) and a small straight metallic head-post was securely cemented to the skull. Animals were given a dose of Carprofen on the day of the surgery, and were allowed to recover for a minimum of 5 days. Prior to the recording session, a small circular craniotomy (1.5 mm of diameter) was opened above the dorsal striatum. During acquisition, animals were head-restrained by a custom-made apparatus and allowed to run on top of a rotatable cylinder. A silicon probe (ASSY 77-H2, Cambridge NeuroTech) was slowly lowered (2-10 *µ*m/s) into the dorsal striatum (∼3 mm from brain surface). We allowed a minimum of 30 minutes before recordings began. Headstage data was digitized at 30 kHz and acquired using the Open-Ephys Acquisition board and Bonsai. In order to align electrophysiology data with pump events, we used the aforementioned “Triggered Control” mode and configured one of the controller’s outputs to report STEP events via TTL, which was then acquired with the same Open-Ephys acquisition board. The syringe pump was positioned 20 cm away from the animal without any electromagnetic shielding material in-between. Protocols were manually triggered by the experimenter. Electrophysiological data shown in (Figure 8) was minimally processed by applying a Butterworth digital bandpass filter (0.3 to 8 kHz). All animal care and experimental procedures were carried out according to the European Directive 2010/63 and pre-approved by the competent authorities, namely, the Champalimaud Animal Welfare Body and the Portuguese Direcção-Geral de Alimentação e Veterinária (DGAV).

## Code and data availability

All code and data related to the device characterization shown in this manuscript are available from https://github.com/fchampalimaud/syringe.pump.manuscript.

## Acknowledgments

We thank Cindy Poo, Hugo Marques and Tiago Monteiro for comments on the manuscript, and Gonçalo Lopes for help with the Bonsai interface.

## Author’s Contribution

B.C. and A.S. collected and analyzed data for the device characterization. P.C. and F.M designed and build the mechanical device. A.S. designed the PCB. D.B. assembled the PCB and initial data collection. L.T. wrote the firmware and graphical user interface, with input from A.S.. S.F. and B.C. designed the animal experiments. S.F. collected and analyzed data from rat behavior experiments. B.C. collected and analyzed data from the acute electrophysical experiments. B.C. made the figures in the manuscript. B.C. and A.S. wrote the manuscript with input from S.F. and L.T.

## Bibliography

Alex Gomez-Marin, Joseph J. Paton, Adam R. Kampff, Rui M. Costa, and Zachary F. Mainen. Big behavioral data: psychology, ethology and the foundations of neuroscience. Nature Neuroscience 2014 17:11, 17:1455–1462, 10 2014. ISSN 1546-1726. doi: 10.1038/nn.3812. URL https://www.nature.com/articles/nn.3812.

F. Nowell Jones and B. F. Skinner. The behavior of organisms: An experimental analysis. The American Journal of Psychology, 52: 659, 10 1939. ISSN 00029556. doi: 10.2307/1416495. URL https://www.jstor.org/stable/1416495?origin=crossref.

Zengcai V. Guo, S. Andrew Hires, Nuo Li, Daniel H. O’Connor, Takaki Komiyama, Eran Ophir, Daniel Huber, Claudia Bonardi, Karin Morandell, Diego Gutnisky, Simon Peron, Ning Long Xu, James Cox, and Karel Svoboda. Procedures for behavioral experiments in head-fixed mice. PLOS ONE, 9:e88678, 2 2014. ISSN 1932-6203. doi: 10.1371/JOURNAL.PONE.0088678. URL https://journals.plos.org/plosone/article?id=10.1371/journal.pone.0088678.

Gonçalo Lopes, Niccolò Bonacchi, João Frazão, Joana P. Neto, Bassam V. Atallah, Sofia Soares, Luís Moreira, Sara Matias, Pavel M. Itskov, Patrícia A. Correia, Roberto E. Medina, Lorenza Calcaterra, Elena Dreosti, Joseph J. Paton, and Adam R. Kampff. Bonsai: an event-based framework for processing and controlling data streams. Frontiers in Neuroinformatics, 9, 4 2015. ISSN 1662-5196. doi: 10.3389/fninf.2015.00007.

Bas Wijnen, Emily J. Hunt, Gerald C. Anzalone, and Joshua M. Pearce. Open-source syringe pump library. PLoS ONE, 9: e107216, 9 2014. ISSN 1932-6203. doi: 10.1371/journal.pone.0107216.

Linda M. Amarante, Jonathan Newport, Meagan Mitchell, Joshua Wilson, and Mark Laubach. An open source syringe pump controller for fluid delivery of multiple volumes. eneuro, 6:ENEURO.0240–19.2019, 9 2019. ISSN 2373-2822. doi: 10.1523/ENEURO.0240-19.2019.

Makoto Ito and Kenji Doya. Validation of decision-making models and analysis of decision variables in the rat basal ganglia. The Journal of Neuroscience, 29:9861–9874, 8 2009. ISSN 0270-6474. doi: 10.1523/JNEUROSCI.6157-08.2009.

Bruno F Cruz, Gonçalo Guiomar, Sofia Soares, Asma Motiwala, Christian K Machens, and Joseph J Paton. Action suppression reveals opponent parallel control via striatal circuits. Nature, 607:521–526, 7 2022.

